# Epidemiological trade-off between intra- and interannual scales in the evolution of aggressiveness in a local plant pathogen population

**DOI:** 10.1101/151068

**Authors:** Frédéric Suffert, Henriette Goyeau, Ivan Sache, Florence Carpentier, Sandrine Gélisse, David Morais, Ghislain Delestre

## Abstract

This preprint has been reviewed and recommended by Peer Community In Evolutionary Biology (http://dx.doi.org/10.24072/pci.evolbiol.100039). The efficiency of plant resistance to fungal pathogen populations is expected to decrease over time, due to its evolution with an increase in the frequency of virulent or highly aggressive strains. This dynamics may differ depending on the scale investigated (annual or pluriannual), particularly for annual crop pathogens with both sexual and asexual reproduction cycles. We assessed this time-scale effect, by comparing aggressiveness changes in a local *Zymoseptoria tritici* population over an eight-month cropping season and a six-year period of wheat monoculture. We collected two pairs of subpopulations to represent the annual and pluriannual scales: from leaf lesions at the beginning and end of a single annual epidemic, and from crop debris at the beginning and end of a six-year period. We assessed two aggressiveness traits – latent period and lesion size – on sympatric and allopatric host varieties. A trend toward decreased latent period concomitant with a significant loss of variability was established during the course of the annual epidemic, but not over the six-year period. Furthermore, a significant cultivar effect (sympatric vs. allopatric) on the average aggressiveness of the isolates revealed host adaptation, arguing that the observed patterns could result from selection. We thus provide an experimental body of evidence of an epidemiological trade-off between the intra- and inter-annual scales in the evolution of aggressiveness in a local plant pathogen population. More aggressive isolates were collected from upper leaves, on which disease severity is usually lower than on the lower part of the plants left in the field as crop debris after harvest. We suggest that these isolates play little role in sexual reproduction, due to an Allee effect (difficulty finding mates at low pathogen densities), particularly as the upper parts of the plant are removed from the field, explaining the lack of transmission of increases in aggressiveness between epidemics.

## INTRODUCTION

Understanding how quickly plant pathogen populations respond to the deployment of host resistance in agrosystems is a real challenge. The role of spatial and temporal variation in host and pathogen life-history traits in the evolutionary trajectories of plant pathogens remains poorly understood (Barrett *et al*., 2008), and we still have few empirical data concerning the ways in which interactions between epidemiological and evolutionary processes influence the generation and maintenance of such variation in host-pathogen interactions (Tack *et al*., 2012). Furthermore, the multidimensional nature of phenotypic adaptation is a key component of evolutionary biology. Common approaches focusing on single traits rather than multiple-trait combinations therefore probably limit our understanding of the adaptive value of a species (Laughlin & Messier, 2015).

Pathogenicity is defined as the ability of a plant pathogen to cause disease, i.e. to damage a plant host. Virulence is the qualitative component of pathogenicity which allows a plant pathogen strain to infect a recognized susceptible host; it is largely shaped by host-pathogen interactions in accordance with the gene-for-gene model (Flor, 1971).

Plant pathologists use the term “aggressiveness” to describe the quantitative variation of pathogenicity on a susceptible host (Shaner *et al*., 1992; Lannou, 2012), which was recently suggested as being conditioned also by minor gene-for-minor gene interactions (Niks *et al*., 2015). Aggressiveness traits are quantitative life-history traits, such as infection efficiency, latent period, sporulation rate, and lesion size (Pariaud *et al*., 2009a).

The efficiency of plant resistance tends to decrease over time, due to the evolution of pathogen populations, with an increase in the frequency of virulent or highly aggressive strains (Geiger & Heun, 1989; Lo *et al*., 2012). A breakdown of qualitative resistance due to a matching increase in pathogen virulence has been observed at different spatiotemporal scales: in a single field after a few infection cycles (Alexander *et al*., 1985; Newton & McGurk, 1991), and in a landscape over several years in response to the deployment of host resistance genes (Hovmøller *et al*., 1993; Kolmer, 2002; Rouxel *et al*., 2003; Goyeau & Lannou, 2011). So-called ‘boom-and-bust’ cycles are typical of rapid selection for particular virulence genes in a pathogen population corresponding to resistance genes present in a local host population (Mundt, 2014). Only a few experiments have addressed the issue of changes in the aggressiveness of pathogen populations over large time scales, due to the complex nature of the relationship between evolution of aggressiveness and evolution of virulence. In the *Puccinia triticina*-*Triticum aestivum* pathosystem, the dominance of a single pathotype is explained not only by its virulence on its sympatric host cultivar, but also by its greater aggressiveness (Goyeau & Lannou, 2011; Pariaud *et al*., 2012). In the wild *Melampsora lini-Linum marginale* pathosystem, a trade-off between the qualitative and quantitative components of pathogenicity may play a key role in generating local adaptation (Thrall & Burdon, 2003). This trade-off may also explain the inconsistency of results for the evolution of aggressiveness following the selection of *Z. tritici* on susceptible vs. moderately resistant wheat cultivars (Ahmed *et al*., 1996; Cowger *et al*., 2002).

The investigation of changes in aggressiveness has paid little attention to the degree of aggressiveness itself, and the most significant changes have been established over the course of an annual epidemic in field experiments with partially resistant cultivars or cultivar mixtures (Newton & McGurk, 1991; Montarry *et al*., 2008; Caffier *et al*., 2016; Delmas *et al*., 2016). In several cases, evolution of aggressiveness was found to be independent of host genetic background or of the virulence genes present in the pathogen population (Andrivon *et al*., 2007; Villaréal & Lannou, 2000). In pea, changes in the aggressiveness of *Didymella pinodes* populations over the course of an annual epidemic differ between winter and spring crops (Laloi *et al.*, 2016), highlighting the potentially complex influences on selection of cropping system, climatic conditions, and epidemiological processes, depending on the nature of the inoculum.

Evolution of aggressiveness due to selection over the course of an annual epidemic may be too weak to be empirically detected, relative to shifts in pathogenicity occurring at larger temporal scales (e.g. Miller *et al*., 1998). Moreover, there may be no selection for aggressiveness traits over small spatiotemporal scales, or this selection may be less intense in natural conditions, for instance due to genetic trade-offs between aggressiveness traits (Laine & Barrès, 2013). Alternatively, selection may be negligible relative to other antagonistic evolutionary forces (e.g., gene flow due to allo-inoculum; McDonald *et al*., 1996; Laloi *et al*., 2016). Selection for greater fitness in pathogen populations is known to be increased by within-host competition among pathogen strains (Zhan & McDonald, 2013), while acknowledging that the more aggressive strains may not always be the more competitive. Co-inoculating a host with a synthetic pathogen population consisting of isolates differing strongly in aggressiveness and then assessing the competition between these isolates is a convenient way to exemplify such a selection (Pariaud *et al*., 2009b; Zhan & McDonald, 2013). Several studies have shown that the adaptation of plant pathogens, in terms of aggressiveness traits, can occur after repeated cycling on the same host. “Serial-passage competition experiments” were designed, with the inoculation of a host plant with a pathogen population, followed by the inoculation of a new set of plants with the offspring of the initial pathogen population, repeated over several cycles: rearing a heterogeneous population of *Puccinia graminis* f. sp. *avenae* separately on two different oats genotypes for seven asexual generations caused the mean infection efficiency of the population to increase by 10–15% by the end of the experiment on the host on which it had been maintained, but not on the other host (Leonard, 1969). In similar “artificial selection experiments”, the use of only the subset of the pathogen population with the highest virulence or aggressiveness to inoculate the next generation of host plants resulted in a shortening of the latent period of an asexual *P. triticina* population after five generations (Lehman & Shaner, 1997).

Some of the epidemiological processes driving selection within a pathogen population can act during the interepidemic period, partly because the local host might change locally (cultivar rotation), so the time-scale to be considered when investigating the evolution of aggressiveness is crucial. However, it is rarely taken into account explicitly: “Most empirical studies have replicated sampling across space rather than through time, based on the argument that assessment across multiple populations in space provides a reasonable surrogate for variation through time” (Tack *et al*., 2012).

Selection for greater aggressiveness during an epidemic period may be followed by reciprocal counter-selection during the subsequent interepidemic period. Greater aggressiveness may impede interepidemic transmission, by limiting the persistence of the host organs on which the pathogen survives (e.g. potato tuber for *P. infestans*; Montarry *et al*., 2007; Pasco *et al*., 2015; Mariette *et al*., 2015) or by decreasing the ability to reproduce sexually (Abang *et al*., 2016; Sommerhalder *et al*., 2011; Suffert *et al*., 2015; 2016). The empirical detection of trade-offs relationship between intra-epidemic multiplication and interepidemic transmission in agrosystems is challenging (Laine & Barrès, 2013): at least two different, nested, selective dynamics act over two different time-scales (annual and pluriannual) under common environmental conditions (same location, same host population) and have yet to be characterized.

The goal of this study was to test the hypothesis of a trade-off relationship between intra-and interepidemic evolutionary dynamics. We therefore investigated changes in the aggressiveness traits of a field population of *Zymoseptoria tritici* at the annual and pluriannual scales. This fungus causes recurrent epidemics of Septoria tritici blotch on wheat, and has a dual sexual-asexual reproduction cycle. At relevant dates characterizing the annual and pluriannual scales, two pairs of *Z. tritici* subpopulations were sampled from a field in which wheat had been grown for several years. The intensity of intra- and interepidemic evolutionary dynamics was investigated by assessing the aggressiveness traits of the fungal isolates *in planta*. We quantified temporal changes in the between-isolate variance within each subpopulation, this variance being expected to decrease with selection. We began by characterizing the epidemiological context in two different ways: we assessed disease variables reflecting the “pathogen pressure” at different dates characterizing key epidemiologic periods, to estimate the temporal continuity in disease dynamics at the annual and pluriannual scales; we also assessed the aggressiveness of the isolates (lesion size and latent period) on both “sympatric” and “allopatric” host cultivars, for isolates sampled early and late over the time course of the experiment, for the detection of local host adaptation patterns.

## MATERIALS AND METHODS

### Host-pathogen system

During the plant-growing season, *Z. tritici* is clonally propagated by asexual pycnidiospores (conidia), which are splash-dispersed upwards, from leaf to leaf. The rate of spread of the epidemic is determined by the number of asexual, embedded infection cycles. Wind-dispersed sexual ascospores, mostly produced on wheat debris during the period between crops, initiates the next epidemic (Suffert *et al*., 2011). Recombination maintains high levels of genetic diversity in *Z. tritici* populations. Selection for both virulence and aggressiveness on wheat cultivars leads to adaptation to the predominant host genotypes (Ahmed *et al*., 1995; 1996; Cowger *et al*., 2002; McDonald *et al*., 1996; Morais *et al*., 2016a).

### Experimental design

A field plot was sown with a wheat cv. Soissons monoculture, by direct drilling without fungicide application, every year from 2007 to 2016, at the Grignon experimental station (Yvelines, France; 48°51'N, 1°58'E), located in the heart of the largest French wheat-producing area (Figure 1). Wheat debris was not removed after harvest and acted as a local source of primary inoculum during the fall (Suffert & Sache, 2011; Morais *et al*., 2016b).

**Figure 1.**
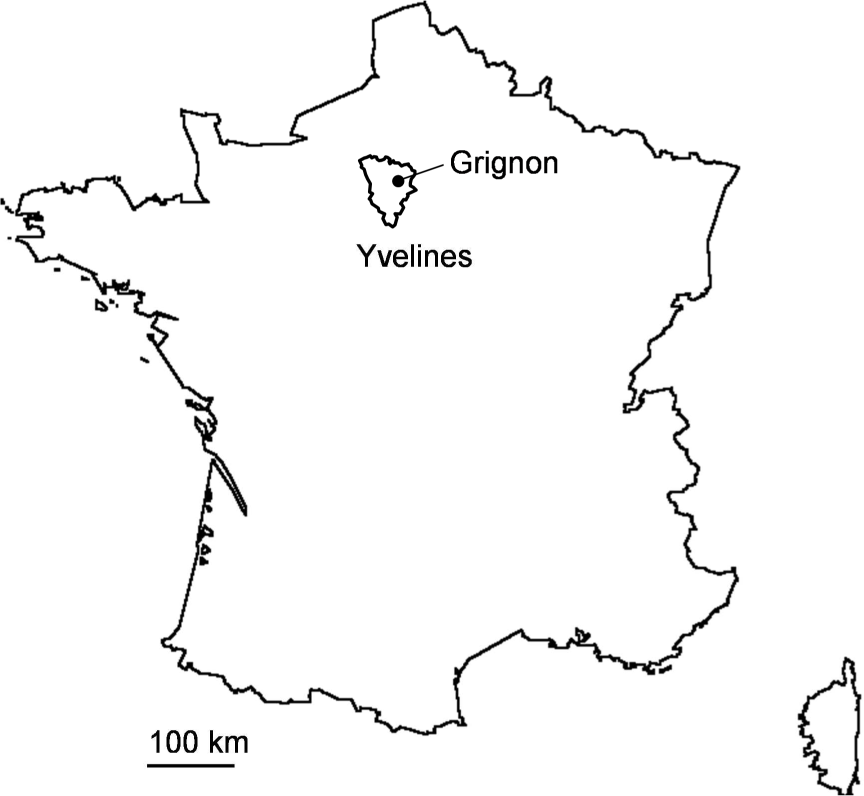
Localization of the study area (Grignon, Yvelines).

Bread wheat cv. Soissons and cv. Apache are both moderately susceptible to Septoria tritici blotch (rated 5 on a 1–10 scale of susceptibility [decreasing from 1 to 10], ARVALIS-Institut du Végétal/CTPS). Resistance is conferred by the *Stb4* and *Stb11* genes in Apache and by an as yet unidentified gene in Soissons (Suffert *et al*., 2013; Thierry Marcel, pers. comm.). Soissons and Apache were among the most popular wheat cultivars in France from 1990 to 2015 and, to a lesser extent, in the area around the study site (Figure 2). Soissons, which was the predominant wheat cultivar in France from 1991 to 2000, subsequently declined in popularity, and accounted for less than 3% of the area under wheat after 2009. Apache, which has been grown in France since 2001, steadily decreased in popularity after 2005; it accounted for 8–12% of the area under wheat in 2009 to 2015. However, Apache remained the most popular cultivar during this period in France generally and in the area surrounding the study site.

**Figure 2.**
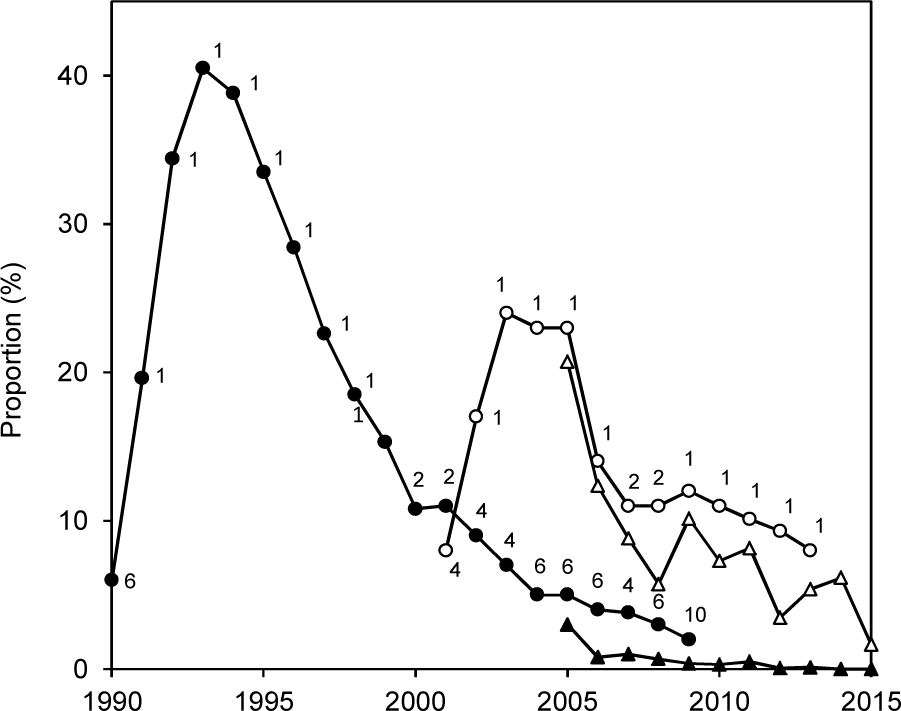
Changes in the proportions of the area under wheat sown with cv. Soissons (black symbols) and cv. Apache (white symbols) from 1990 to 2015 in France (round symbols) and from 2005 to 2015 in the Yvelines (trianglular symbols). The numbers correspond to the ranking of each cultivar in France (1 = the most deployed, 2 = the second most deployed, etc.; data FranceAgriMer).

In our study, Soissons was referred to as “sympatric” host cultivar for the tested pathogen isolates because they were directly sourced from it. As a once-predominant cultivar now in decline, Soissons was assumed to have played a major role in the overall evolutionary trajectory of pathogen populations in France. Soissons played a specific role in the experimental study area because it was grown there in monoculture for eight years. Apache was considered to be an “allopatric” host cultivar. It partially replaced Soissons as the predominant cultivar in France, but probably played a less important role than Soissons in the evolutionary trajectory of the local pathogen population, with potentially some isolates immigrating from commercial fields located around the field plot. We therefore considered the most likely origins of the local pathogen subpopulations to be, firstly, cv. Soissons, and, secondly, cv. Apache.

### Disease dynamics over ten years

The temporal dynamics of pathogen pressure in the wheat monoculture plot was characterized from 2008 to 2016 with five quantitative disease variables assessed in field conditions: the amount of primary inoculum at the onset of the epidemic, the earliness of the attack, winter disease severity at the late tillering stage, spring disease severity during the stem extension stage, and spring disease severity during the grain-filling period (see complete definitions in Figure 3). The overall continuity in disease dynamics was investigated: i) at the intra-epidemic scale, by assessing correlations between variables recorded during a single annual epidemic; and ii) at the interepidemic scale, by assessing correlations between variables (the same or different) recorded during two successive epidemics.

**Figure 3.**
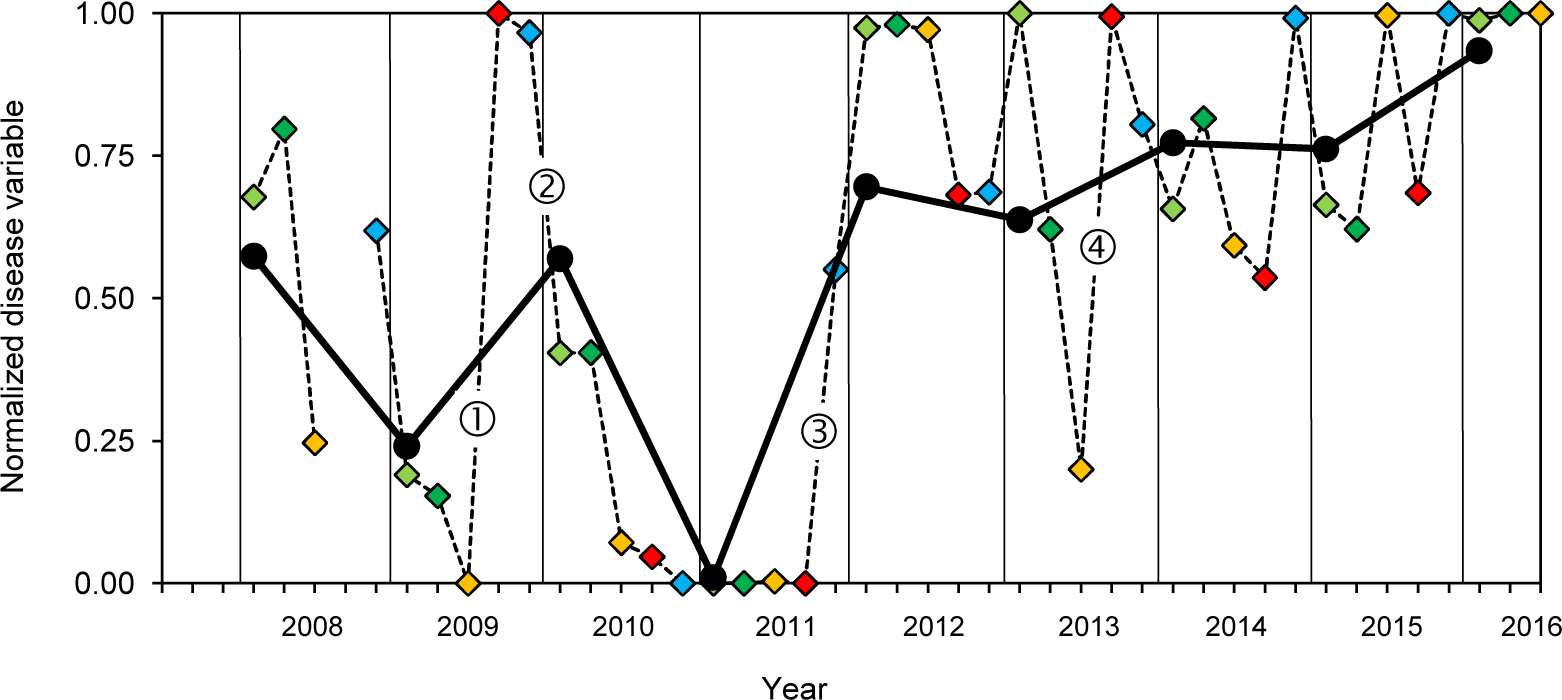
Normalized disease variables used to quantify *Zymoseptoria tritici*pathogen pressure in the wheat monoculture plot from 2008 to 2016. Each variable was normalized to give a value in the range 0–1 (from the lowest to the highest annual value calculated over the 2008–2016 period). An overall index (black circles), calculated as the mean of the five variables for each epidemic period, was used as a proxy for annual pathogen pressure. The “earliness of attack” (blue diamonds) corresponds to the date on which the epidemic reached a threshold intensity (mean date on which the proportion of diseased plants reached 5% and the date on which the mean disease severity for leaf layer L1 reached 20%); 0 = earliest date, 1 = latest date. “Winter disease severity” at the late tillering stage (light green diamonds) corresponds to the mean disease severity assessed on leaf layer L4 from mid-January to mid-February. “Early-spring disease severity” during stem extension (dark green diamonds) corresponds to the mean disease severity assessed on leaf layers L5 and L6 from mid-March to mid-April. “Late-spring disease severity” during the grain-filling period (yellow diamonds) corresponds to the mean disease severity assessed on leaf layer F2 in early June. The “amount of primary inoculum” available at the end of the epidemic period (red diamonds) corresponds to the mean number of ascospores collected from 1 g of wheat debris (mean number of ascospores collected in mid-October and in mid-November; 0 = lowest primary inoculum pressure, 1 = highest inoculum pressure). The circled numbers corresponds to discontinuities in pathogen pressure (change of at least 0.5 on a standardized scale, i.e. half the maximal amplitude observed during the 2008–2016 period for each variable normalized in the range 0–1) between two disease variables successive in time and positively correlated during the 2008–2016 period.

### Sampling of pathogen subpopulations

Two pairs of pathogen subpopulations were sampled from the monoculture field plot (Figure 4). The first pair (2 × 15 isolates; Table S1): the “initial ascospore-derived subpopulation” (Ai-2009) and the “final ascospore-derived subpopulation” (Af-2015), was obtained from ascospores ejected from infected debris during the 2009 and 2015 interepidemic periods, respectively (on 9 October 2009 and from 7 September 2015 to 13 October 2015, respectively; Morais *et al.*, 2016b). The selective trajectory of these two subpopulations was affected by several reproduction cycles on wheat cv. Soissons cultivated for two and seven years, respectively (Figure 4). The second pair (2 × 15 isolates; Table S1): “initial conidial subpopulation” (Ci-2009) and “final conidial subpopulation” (Cf-2010), was obtained from leaf lesions during the 2009–2010 epidemic period. Ci-2009 isolates were collected at the beginning of the epidemic (from 24 November to 8 December 2009), from the first 15 lesions detected on seedlings. Most isolates belonging to this subpopulation would have come from debris present in the monoculture field plot (Morais *et al*., 2016b). Cf-2010 isolates were collected at the end of the same epidemic (on 12 July 2010), from 15 lesions on the antepenultimate (F3) or penultimate leaf (F2). The vertical disease profile suggests that most of these lesions were caused by secondary reinfection with local inoculum rather than the arrival of immigrant isolates from contaminated debris in distant fields (Suffert & Sache, 2011).

**Figure 4.**
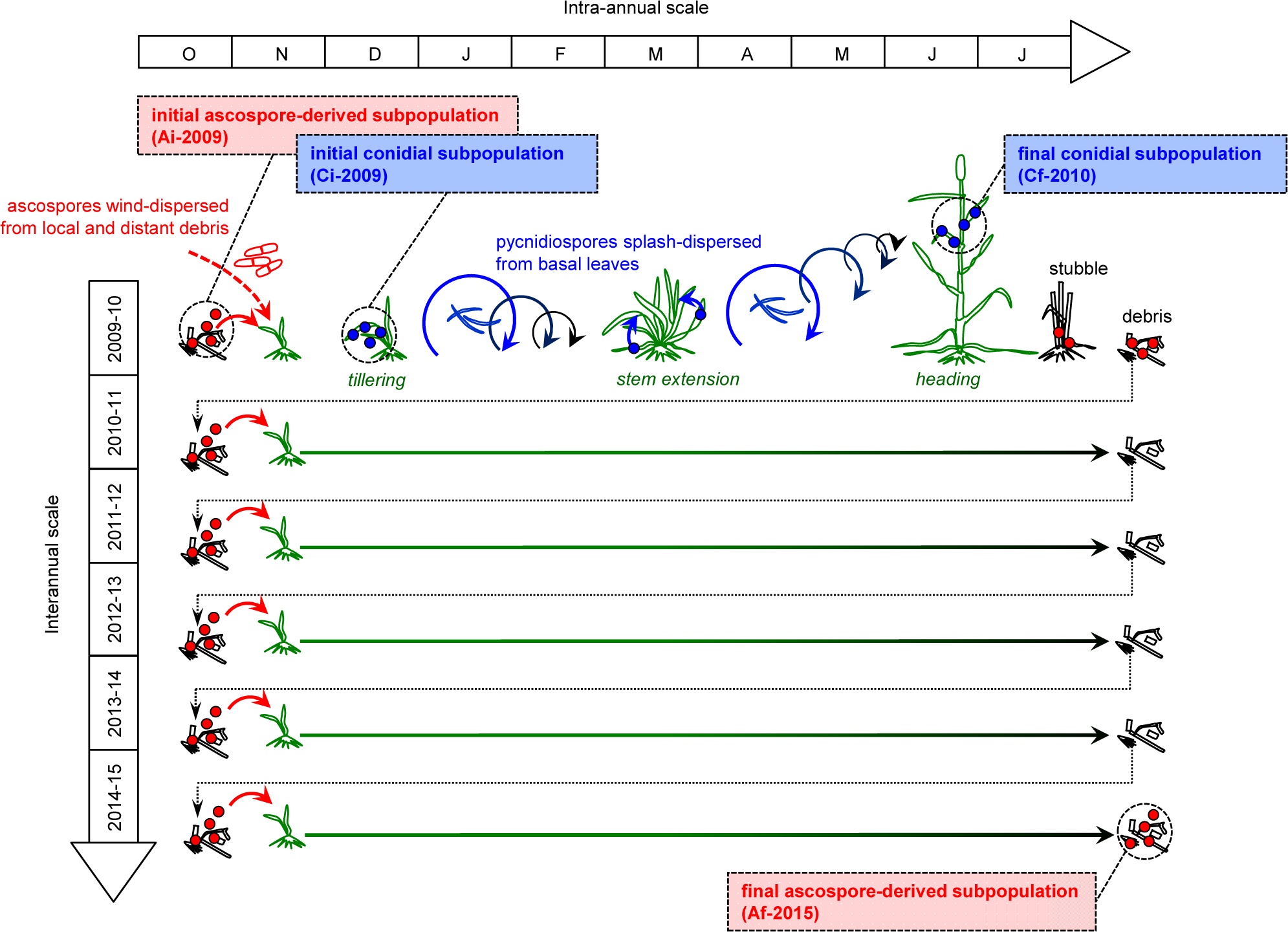
Dynamics of two pairs of *Zymoseptoria tritici*subpopulations (2 × 2 × 15 isolates) collected in the wheat monoculture plot from infected leaves (initial and final conidial subpopulations in 2009–2010) or from debris (initial and final ascospore-derived subpopulations in 2009 and 2015, respectively).

### Assessment of aggressiveness traits

The aggressiveness of the 60 isolates was assessed on adult plants, with the method developed by Suffert *et al.* (2013), in two independent greenhouse trials. Ci-2009 and Cf-2010 isolates were tested in 2011, on F1 and F2 leaves (2 × 4 replicates) of cv. Soissons. Ai-2009 and Af-2015 isolates were tested in 2015, on F1 leaves (6 replicates) of both cv. Soissons and cv. Apache.

Wheat seedlings were vernalized in a cold growth chamber for seven weeks, then brought to the greenhouse at 12–20°C and transplanted into larger pots. Three stems per plant were kept. The thermal time *t*, expressed in degree-days post inoculation (ddpi), was calculated, starting from the inoculation date, by summing the daily mean air temperature, using a base temperature of 0°C. For each isolate, subcultures were prepared on Petri dishes. Aqueous conidial suspensions were adjusted to a final concentration of 10^5^ conidia.mL^-1^ and applied with a paintbrush along a 25 mm-long section of the adaxial face of the leaves of the main tiller of each plant. Leaves were then enclosed for 72 h in a slightly damp transparent bag to promote infection. Lesion size, estimated by determining the sporulating area, was assessed by eye on each leaf with a hand lens, as the percentage of the inoculated leaf surface bearing pycnidia. Assessments were performed twice weekly, from inoculation to leaf senescence (11 and 10 assessments in 2011 and 2015, respectively). A Gompertz growth curve was fitted to the values of lesion size recorded over time on each leaf, as described by Suffert *et al*. (2013) using the S-PLUS 6.0 software (Lucent Technologies, Inc.). The latent period, defined here as the time elapsed from inoculation to 5% of the maximum percentage of the area bearing pycnidia, was calculated from the parameters of the Gompertz growth curve estimated for each leaf and is expressed in ddpi.

### Data analysis

The aggressiveness traits of each pair of pathogen subpopulations were assessed in a separate greenhouse trial, by two different people. This assessment is assessor-dependent and influenced by environmental conditions, so data from the two trials could not be pooled. A nested ANOVA was used to assess differences between the two subpopulations of each pair, for each aggressiveness trait (maximum lesion size and latent period). For the pair of conidial subpopulations, we considered subpopulation (initial Ci-2009, final Cf-2010) and cultivar (Soissons, Apache) as fixed effects, isolates as a random factor nested within subpopulation, and their interactions. For the pair of ascospore-derived subpopulations, we considered subpopulation (initial Ai-2009, final Af-2015) and leaf layer as fixed effects, isolate as a random factor nested within subpopulation, and their interactions. We determined whether the between-subpopulation variance of lesion size and latent period on cv. Soissons decreased significantly between Ci-2009 and Cf-2010 and between Ai-2009 and Af-2015, by calculating the null distribution of the ratio of variances in permutation tests (100,000 permutations). ANOVA were performed using the S-PLUS 6.0 software (Lucent Technologies, Inc.) and permutation tests using the R software.

## RESULTS

### Correlation between disease variables at different time-scales

The correlation between the earliness of attack and the amount of primary inoculum available at the end of the previous epidemic period was clearly positive but not statistically significant (ρ = 0.505 with *p* > 0.1; Table 1): the higher the number of ascospores discharged from wheat debris in the fall, the earlier the first symptoms appeared after seedling emergence, consistent with previous experimental results (Suffert & Sache, 2011). The correlation between earliness of attack and disease severity in the current season was positive, whatever the period of severity assessment (ρ= 0.407 with *p*> 0.1 for winter disease severity; ρ= 0.491 with *p*> 0.1 for early-spring disease severity; ρ= 0.663 with *p*< 0.1 for late-spring disease severity): the earlier the onset of the first symptoms, the higher disease severity. The correlation between disease severity assessed on two dates within the same epidemic period was highly positive and statistically significant (ρ= 0.761 with *p*< 0.05 and ρ= 0.831 with *p*< 0.01), consistent with the generally accepted view that Septoria tritici blotch severity is proportional to the intensity of secondary infections, driven by pycnidiopores splash-dispersed from existing, sporulating lesions.

**Table 1.** Intra- and interannual Spearman's rank correlation coefficients (ρ) for the relationships between five disease variables (see definitions in Figure 3) characterizing the pathogen pressure in the wheat monoculture plot from 2008 to 2016. Correlations were assessed for the same and different variables from two successive annual epidemics, and for different variables from the same annual epidemic. Ávalues higher than 0.50 appears in bold and significant ρ values are indicated by *** (*p*< 0.01), ** (*p*< 0.05) and * (*p*< 0.1).

| Variable of epidemic year n - 1 | Variable of epidemic year n |  |  |  |  |
| --- | --- | --- | --- | --- | --- |
|  | Earliness of attack | Winter disease severity | Early-spring disease severity | Late-spring disease severity | Amount of primary inoculum<sup>1</sup> |
| Earliness of attack | 0.430 | -0.270 | -0.018 | 0.180 | -0.609 |
| Winter disease severity | 0.204 | 0.211 | 0.042 | -0.024 | 0.342 |
| Early-spring disease severity | 0.299 | 0.295 | -0.084 | 0.103 | **0.559** |
| Late-spring disease severity | **0.527** | **0.518** | 0.301 | 0.400 | **0.609** |
| Amount of primary inoculum<sup>1</sup> | **0.505** | -0.036 | -0.082 | -0.173 | -0.145 |
| Variable of epidemic year n |  |  |  |  |  |
| Earliness of attack |  | 0.407 | 0.491 | **0.663 \*** | 0.054 |
| Winter disease severity |  |  | **0.761 \*\*** | **0.620 \*** | 0.427 |
| Early-spring disease severity |  |  |  | **0.831 \*\*\*** | 0.209 |
| Late-spring disease severity |  |  |  |  | 0.127 |
| Amount of primary inoculum<sup>1</sup> |  |  |  |  |  |

<sup>1</sup> available for the subsequent epidemic period

The correlation between disease severity and the amount of primary inoculum available for the subsequent epidemic period was positive but not statistically significant (*p* > 0.1). Higher correlation coefficients were obtained for earlier assessments of disease severity during the epidemic period (ρ = 0.427 for winter disease severity; ρ = 0.209 for early-spring severity; ρ = 0.127 for late-spring disease severity; Table 1). This is consistent with the hypothesis that sexual reproduction is a density-dependent process (positively correlated with the density of lesions, i.e. disease severity; Suffert, unpubl. data) probably occurring towards the base of the plants, where the proportion of mature *Z. tritici* pseudothecia among the overall fruiting bodies (pycnidia and pseudothecia) is systematically higher than on the upper part of plants over the course of an epidemic (Eriksen & Munk, 2003). This finding is also supported by previous experimental results showing that ascospore production is generally greatest after the most severe previous epidemics (Cowger *et al*., 2002).

The distribution of the coefficients of correlation between different disease variables from a single annual epidemic was compared with that from two successive annual epidemics (Figure 5). Only 5/25 coefficients of correlation between different disease variables from two successive annual epidemics (interepidemic scale) exceeded 0.5, versus 4/10 for variables from a single annual epidemic (intra-epidemic scale); no coefficients of correlation between different disease variables from a single annual epidemic were negative, versus 7/25 for variables from two successive annual epidemics. Because of the low proportion of significant coefficients of correlation, no statistical test was suitable for comparing the two distributions. The correlation between the same disease variables from two successive annual epidemics was low (-0.145 ≤ ρ≤ 0.429; Table 1). The overall temporal continuity of pathogen pressure was tighter between successive epidemiological stages than between identical epidemiological stages from two successive annual epidemics.

**Figure 5.**
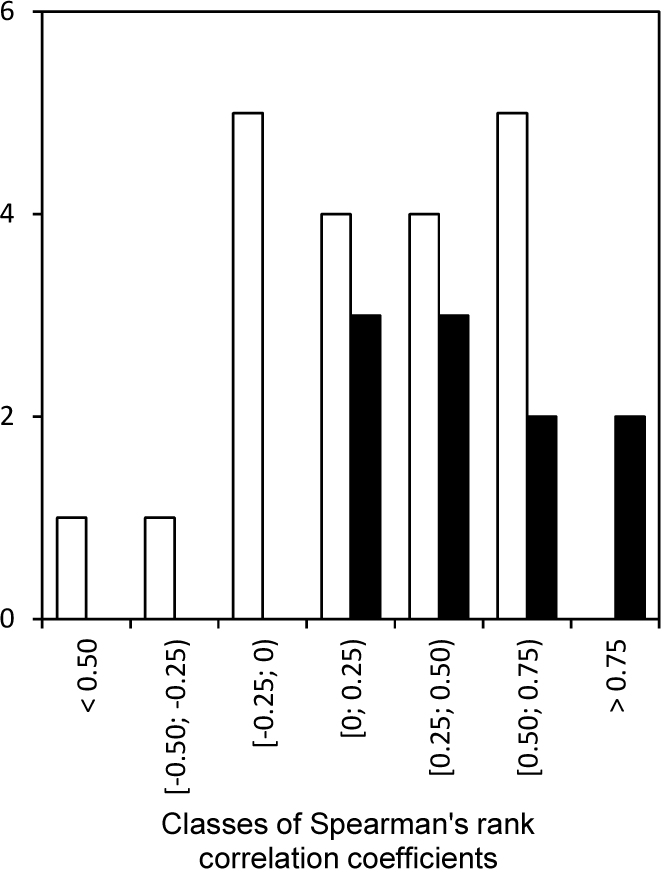
Distribution of the coefficients of correlation between different disease variables from a single annual epidemic (black bars; intra-epidemic scale) and between different disease variables from two successive annual epidemics (white bars; interepidemic scale).

We defined a “significant discontinuity” in pathogen pressure as a change of at least 0.5 on a standardized scale between two disease variables successive in time and positively correlated during the 2008–2016 period, i.e. between i) amount of primary inoculum and earliness of attack; ii) earliness of attack and winter disease severity; iii) winter disease severity and early-spring disease severity; iv) early-spring disease severity and late-spring disease severity; and v) winter disease severity and the amount of primary inoculum at the beginning of the subsequent epidemic (Figure 3). No major discontinuity was identified from late-spring 2010 to fall 2011 (very low pathogen pressure following exceptionally low rainfall levels during the spring of 2010). Only three significant discontinuities were identified: i) during the 2009 interepidemic period, in the fall, with a large amount of primary inoculum despite moderate disease severity in the previous spring (point ① in Figure 3); ii) during the 2009–2010 spring intra-epidemic period, with an early attack followed by moderate winter disease severity, due to weather conditions not conducive to disease development (low rainfall, low temperature; point ② in Figure 3); iii) during the 2011–2012 late-winter intra-epidemic period, with very small amounts of primary inoculum followed by a very high winter disease severity, due to weather conditions conducive to disease development (high rainfall and temperature; point ③ in Figure 3). The discontinuity between late-spring disease severity in 2013 and the amount of primary inoculum at the beginning of the subsequent epidemic (point ④ in Figure 3) was not considered “significant” as previously defined, because the two disease variables were not positively correlated during the 2008–2016 period (Table 1). From 2012 to 2016, pathogen pressure remained high and no significant discontinuity was found.

### Dynamics of lesion development

Of the 240 and 360 lesion development curves obtained in the 2011 and 2015 greenhouse trials, 15 and 43, respectively, were excluded from the statistical analysis due to either a partial failure of inoculation or a lack of model convergence. An example of the data sets obtained for two isolates, one from Cf-2010 and one from Af-2015, is provided in Figure 6, highlighting the intra-isolate variance of lesion size obtained after fitting. Figures 7a and 7c illustrate the mean development of lesions for each isolate-cultivar combination; each curve represents the mean growth of lesions calculated with six or eight replicates, for conidial (7a) and ascospore-derived (7c) subpopulations, respectively. Figures 7b and 7d illustrate the mean development of lesions for each subpopulation × cultivar interaction; each curve represents the average growth of lesions induced by the 15 isolates of each subpopulation. The disease progress curve of Cf-2010 appears to be slightly ahead of that for Ci-2009, suggesting that the final conidial subpopulation was more aggressive than the initial conidial subpopulation. The disease progress curves of Af-2015 and Ai-2009 overlapped, suggesting an absence of difference in aggressiveness between the two ascospore-derived subpopulations. Moreover, the disease progress curves of subpopulations tested on Apache showed a marked delay relative to the curves obtained for Soissons, suggesting that Ai-2009 and Af-2015 were less adapted to their allopatric host.

**Figure 6.**
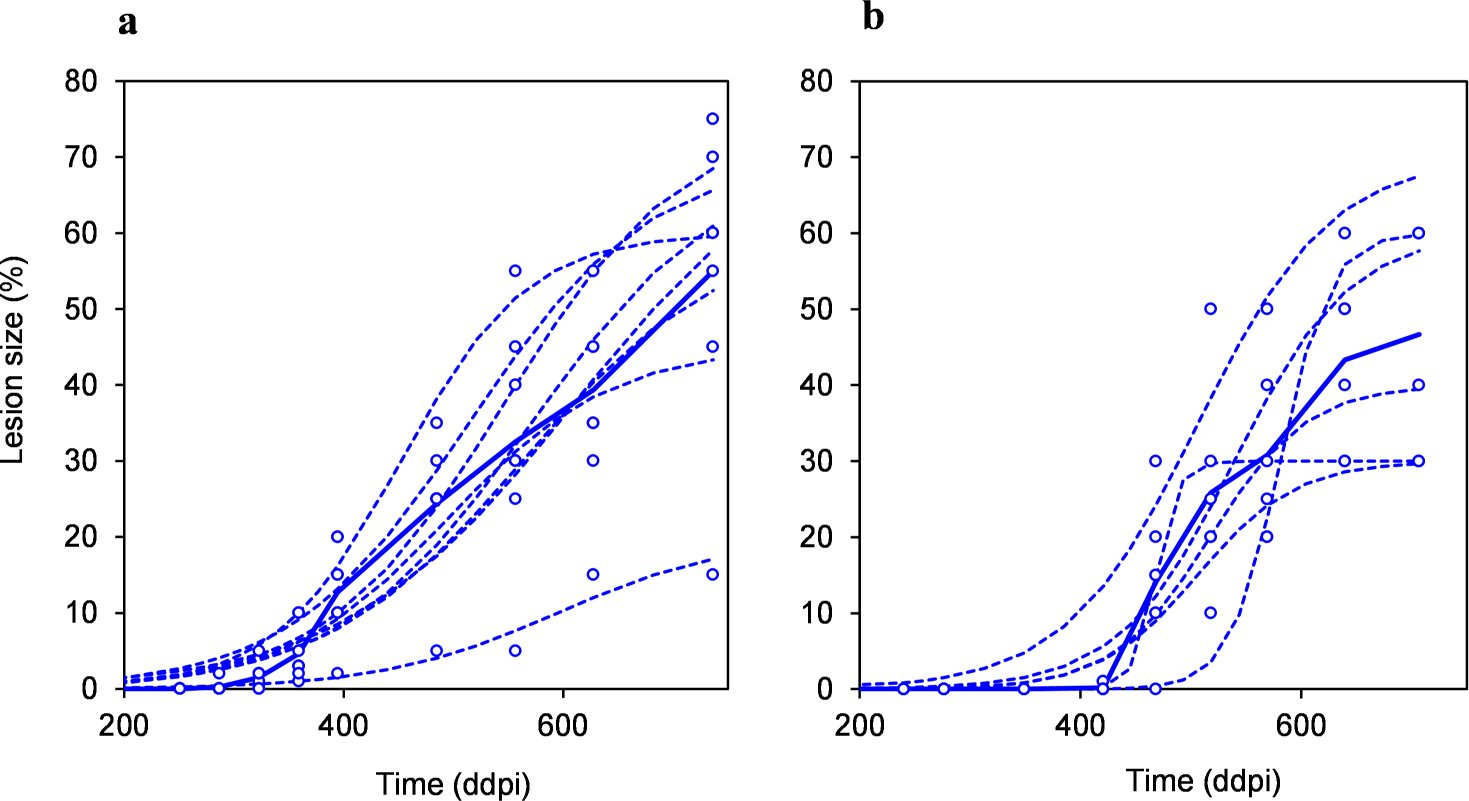
Growth of lesions on cv. Soissons caused by Zymoseptoria tritici isolates I21 (a) and I47 (b), illustrating the behavior of the different pathogen subpopulations. Lesion size (sporulating area) is expressed as a % of the inoculated area; time is expressed in degree-days post inoculation (ddpi) since the sowing date (base temperature 0 °C). The dotted curves correspond to the model fitted to the experimental data for each replicate (four F1 and four F2 leaves for I21; six F1 leaves for I47); the bold curve corresponds to the mean growth of the lesions induced by each isolate.

**Figure 7.**
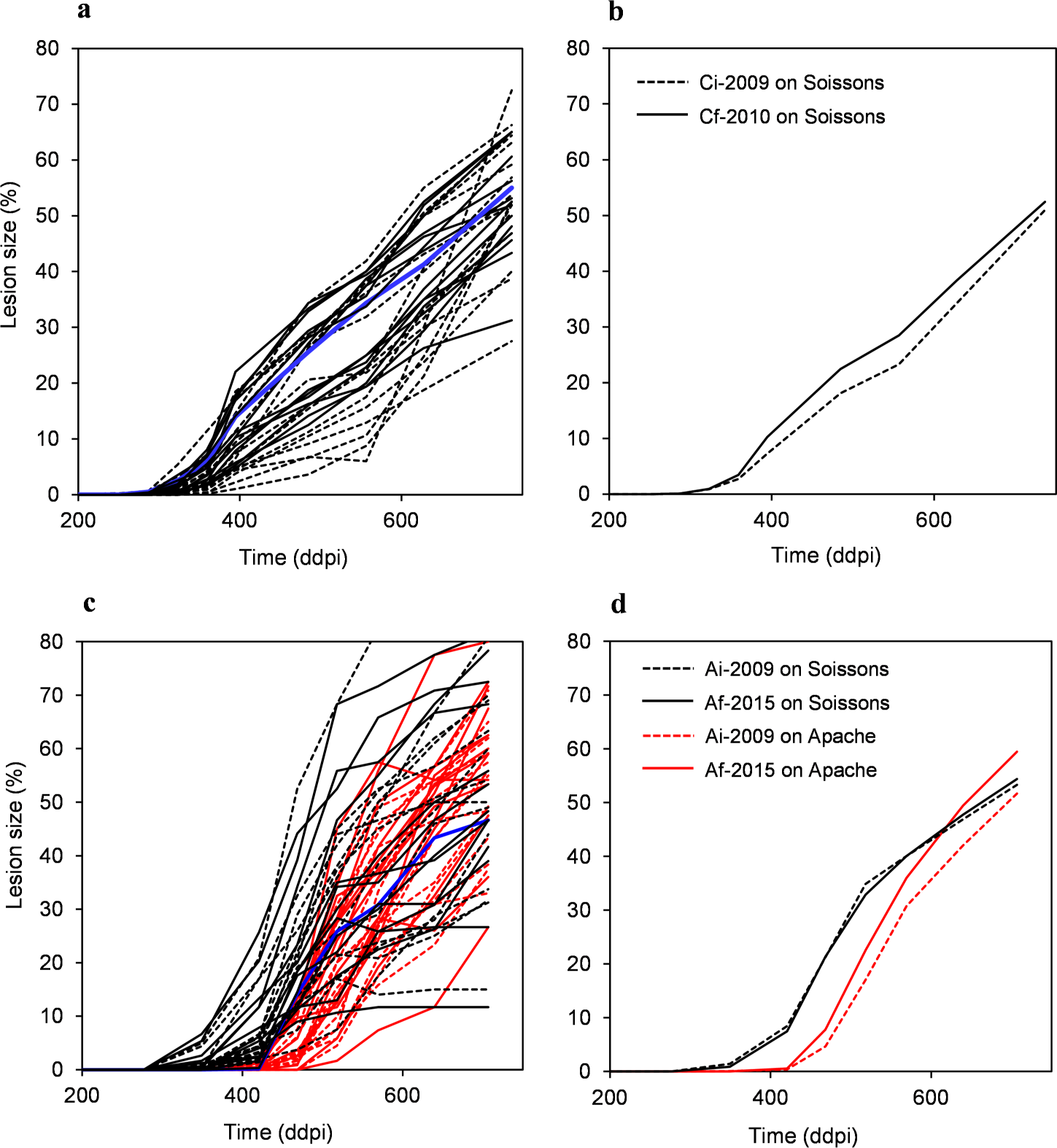
Growth of lesions caused by two pairs of *Zymoseptoria tritici*subpopulations (60 isolates) collected from infected leaves (initial [Ci-2009] and final [Cf-2010] conidial subpopulations in 2009–2010) or from debris (initial [Ai-2009] and final [Af-2015] ascospore-derived subpopulations in 2009 and 2015, respectively) in the wheat monoculture plot. Each curve in 7a (2 × 15) and 7c (4 × 15) shows the mean growth of lesions induced by a given isolate on cv. Soissons (blue lines correspond to isolates I21 [7a] and I47 [7c]; see Figure 6). Each curve in 7b and 7d shows the mean growth of lesions induced by the 15 isolates of each subpopulation (initial populations shown as dotted lines and final populations as bold lines) on cv. Soissons (black lines) and Apache (red lines).

### Subpopulation and cultivar effects on aggressiveness traits

Between-isolate variability was high for both latent period and lesion size, revealing a high level of phenotypic diversity in the four pathogen subpopulations: in the conidial and ascospore-derived subpopulations, lesion size (*p*= 0.052 and *p* < 0.001, respectively) and latent period (*p*< 0.001) differed significantly between isolates (Table 2 and 3).

**Table 2.** Analysis of variance for latent period and lesion size of the initial (Ci-2009) and final (Cf-2010) conidial *Zymoseptoria tritici*subpopulations

| Source of variance <sup>a</sup> | df | Latent period |  |  | Lesion size |  |  |
| --- | --- | --- | --- | --- | --- | --- | --- |
|  |  | MS | F | p | MS | F | p |
| Population P | 1 | 22036 | 2.9 | **0.087** | 10 | 0.02 | 0.881 |
| Isolate I(P) | 28 | 6990 | 4.5 | **<0.001** | 693 | 1.5 | **0.052** |
| Leaf layer L | 1 | 19031 | 12.2 | **0.002** | 5357 | 11.9 | **<0.001** |
| Residual |  | 1562 |  |  | 483 |  |  |

<sup>a</sup> The P×L and I(P)×L interactions were non-significant.

**Table 3.**
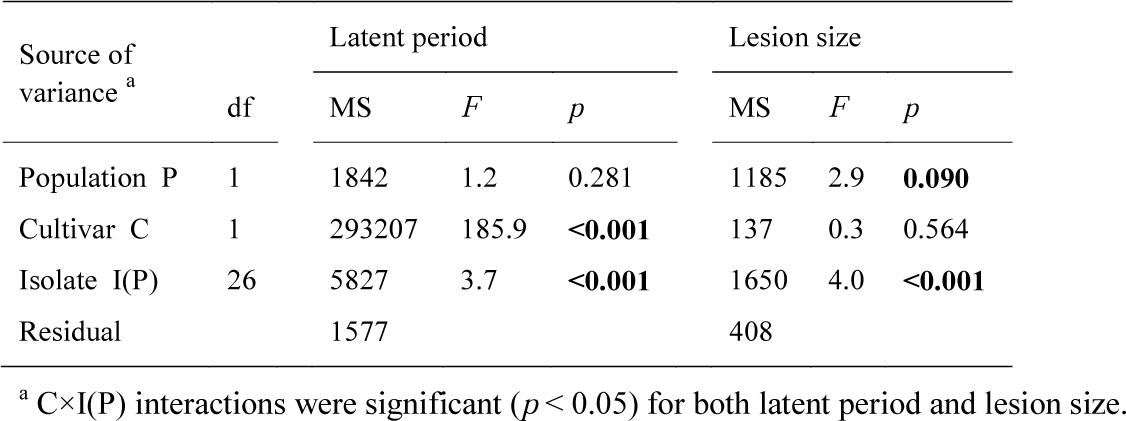
Analysis of variance for latent period and lesion size of the initial (Ai-2009) and final (Af-2015) ascospore-derived *Zymoseptoria tritici*subpopulations

No difference in between-isolate variance (Vg) was detected between Ci-2009 and Cf-2010 for lesion size (Vg(Ci-2009)/Vg(Cf-2010) = 1.343; *p* = 0.608), whereas Vg significantly decreased over time for latent period (Vg(Ci-2009)/Vg(Cf-2010) = 3.835; *p* = 0.042). No difference in between-isolate variance was detected between Ai-2009 and Af-2015 for either lesion size (Vg(Ai-2009)/Vg(Af-2015) = 1.289; p = 0.643) or latent period (Vg(Ai-2009)/Vg(Af-2015) = 0.677; *p* = 0.442). Leaf layer was also a source of considerable variation for both lesion size and latent period (*p* < 0.001 and *p* = 0.002, respectively; Table 2).

Latent period was shorter in Cf-2010 than in Ci-2009 (428.0 ddpi vs. 409.1 ddpi; data not shown), although the difference cannot be consider as statistically significant (*p* = 0.087; Table 2); there was not difference in lesion size between these two subpopulations (*p*= 0.881).

Lesion size was larger in Af-2015 than in Ai-2009 (61.9 % vs. 58.0 %), although the difference cannot be consider as statistically significant too (*p* = 0.090; Table 3). No significant difference in latent period was detected (*p* = 0.281). The cultivar effect was significant only for latent period, which was longer in the allopatric cultivar than in the sympatric cultivar (491.4 ddpi in Apache vs. 426.3 ddpi in Soissons; *p* < 0.001; Table 3). The aggressiveness of isolates depended on the cultivar on which it was assessed. An ANOVA considering each cultivar (Soissons, Apache) separately identified the origin of the population effect: on Soissons, there was no significant difference in aggressiveness between Ai-2009 and Af-2015, for either lesion size (59.1 % for 2009 vs. 59.5 % for 2015; *p* = 0.863; Table 4) or latent period (427.9 ddpi for 2009 vs. 424.8 ddpi for 2015; *p* = 0.654), contrary to Ci-2009 and Cf-2010; on Apache, the population effect was significant only for lesion size, which was larger for Af-2015 than for Ai-2009 (64.2 % vs. 57.1 %; *p* = 0.037).

**Table 4.** Analyses of variance for latent period and lesion size of the initial (Ai-2009) and final (Af-2015) ascospore-derived *Zymoseptoria tritici*subpopulations performed independently for the wheat cv. Soissons and cv. Apache.

|  | Source of variance | df | Latent period |  |  | Lesion size |  |  |
| --- | --- | --- | --- | --- | --- | --- | --- | --- |
|  |  |  | MS | *F* | *p* | MS | *F* | *p* |
| Soissons |  |  |  |  |  |  |  |  |
|  | Population P | 1 | 306 | 0.2 | 0.654 | 7 | 0.03 | 0.863 |
|  | Isolate I(P) | 28 | 4228 | 2.8 | **<0.001** | 1788 | 7.7 | **<0.001** |
|  | Residual |  | 1510 |  |  | 233 |  |  |
| Apache |  |  |  |  |  |  |  |  |
|  | Population P | 1 | 558 | 0.4 | 0.545 | 2050 | 4.5 | **0.037** |
|  | Isolate I(P) | 28 | 4274 | 2.8 | **<0.001** | 670 | 1.5 | **0.099** |
|  | Residual |  | 1510 |  |  | 456 |  |  |

## DISCUSSION

The correlation between disease variables assessed at different epidemiological time-scales revealed intra- and interannual continuity in pathogen pressure, consistent with current knowledge concerning the epidemiology of Septoria tritici blotch. The three significant discontinuities identified during the six-year period may have been due to environmental factors, such as weather conditions in particular.

The significant effect of cultivar on the latent period of one of the two pairs of *Z. tritici* subpopulations may reflect the difference in resistance between the two cultivars. This assumption is however invalidated by results obtained in similar experimental conditions (same study area, same cultivars tested; Morais *et al*., 2016a). Rather, it may indicate local host adaptation (better performance on the “local” vs. “foreign” host): considering the latent period, after several years of monoculture the resident pathogen population becomes more adapted to sympatric host. This is consistent with the differential adaptation of resident and immigrant *Z. tritici* subpopulations to wheat cultivars previously established in the same study area (Morais *et al*., 2016a) and elsewhere for the same pathosystem (Ahmed *et al.*; 1995, 1996), and, more generally, with evolutionary concepts (Kaltz & Shykoff, 1998; Gandon & Van Zandt, 1998; Kawecki & Ebert, 2004; Tack *et al*., 2012). Considering the lesion size, this conclusion must be however qualified by the gain of adaptation observed to the allopatric host (only) over the six-year period, which can be consider as an evidence of maladaptation (worse performance on the “local” vs. “foreign” host; Kaltz *et al*. 1999). This result is also consistent with those obtained by Morais *et al*. (2016a) from the comparison of the latent period of the immigrant and resident subpopulations on the two cultivars. A host shift alters the current selection environment in which resident strains underwent preadaptation, decreasing competitive ability; resident strains may therefore perform poorly on their new host. Several studies have shown that virulence and aggressiveness of *Z. tritici* populations increase over time on predominant host genotypes, regardless of the type of host resistance in the plant on which they originated (Zhan & McDonald, 2013). However, the results of Ahmed *et al*. (1996) and Cowger *et al*. (2002) concerning selection for higher levels of aggressiveness on susceptible vs. moderately resistant wheat cultivars were inconsistent, possibly caused by an artifact due to a genetic trade-off between virulence and aggressiveness (Zhan *et al*., 2002).

Finally, the overall temporal continuity in disease development over the six-year period and the evidence of local host adaptation provide two arguments supporting that the increase in aggressiveness at the annual scale and the stability of aggressiveness at the pluriannual scale, are due – at least partly – to deterministic evolutionary forces (as opposed to chance), such as selection acting on quantitative traits.

In our experiment, the selective processes driving epidemics at the intra- and interannual scales had antagonistic consequences for the long-term evolution of aggressiveness in the local pathogen population.

A difference in aggressiveness was observed between the initial and final conidial subpopulations (Ci-2009, Cf-2010) expressed on the sympatric host cv. Soissons over the annual time-scale. The dynamics of lesion development were different. On average, the latent period was shorter for the final conidial subpopulation than for the initial one. The difference, however, was not statistically significant at *p* = 0.05 (*p* = 0.087). While there was no difference in lesion size when assessed on adult plants at 18.1°C (spring conditions), lesion size was larger for the isolates of the final conidial subpopulation than for those of the initial conidial subpopulation when assessed on seedlings at 8.9°C (winter conditions) (Suffert *et al*., 2015). These results suggest that aggressiveness increased over the course of a single annual epidemic, although we could not demonstrate it formally. The trend observed during a single year results from differential selection effects that would have needed to be maintained during several years to reveal more significant effects. This evolution reflects a pattern of adaptation, interpreted as the outcome of short-term selection driven by seasonal environmental conditions. Our interpretation is supported by the significant decrease in the between-isolate variance for latent period at the intra-annual scale, compared to the stability of this variance at the interannual scale. McDonald *et al.* (1996) have already suggested that selection affects the genetic structure of *Z. tritici* populations, but they found no experimental evidence for adaptation to any of the host genotypes over the growing season. However, the neutral genetic markers used in their study were not appropriate; the use of markers of aggressiveness or the phenotyping of isolates would have been more relevant approaches. Moreover, sample size was too small to allow the detection a change in the frequency of pathogen genotypes: the probability of detecting the same clone several times is very low with respect to the high population diversity. Finally, with the experimental design used by McDonald *et al.* (1996), it was not possible to exclude the possibility of sexual reproduction during the growing season (Chen & wMcDonald *et al*, 1996; Duvivier *et al.*, 2015), which might conceal the effects of short-term selection.

By contrast, no difference in aggressiveness (latent period or lesion size) was found between the initial and final ascospore-derived subpopulations (Ai-2009, Af-2015) expressed on the sympatric host cv. Soissons for analyses over the pluriannual time-scale. Lesion dynamics were identical and there were no significant differences in curve parameters. There was, therefore, no increase in the aggressiveness of the local pathogen population after several years of wheat monoculture, and no selection effect was detectable. The absence of change from 2009 to 2015 may have been concealed by the impact of immigrant strains (clouds of ascospores released from distant debris) exhibiting an evolutionary trajectory different from that of the resident strains. A previous comparison of the aggressiveness of resident and immigrant strains on adult plants, as in this study, however, demonstrated that most of the primary inoculum originated within the field (Morais *et al.,* 2016a).

Selection led to a short-term increase in the aggressiveness of the pathogen subpopulation primarily responsible for secondary infections (Suffert *et al*., 2015) at the annual scale, with no significant impact at the pluriannual scale. Highly aggressive strains could be selected in the pathogen population after only a few embedded asexual multiplication cycles because sexual reproduction plays a lesser role in disease development during the intra-epidemic period than during the interepidemic period. The impact of such intra-annual selection was nullified at the beginning of the next epidemic, probably because sexual reproduction played a crucial role during the early epidemic stages, in which ascospores are the main form of primary inoculum (Morais *et al*., 2016b; Suffert & Sache, 2011).

Using a field design connecting epidemic and interepidemic periods, we provide experimental evidence of an epidemiological trade-off between the intra- and interannual scales in the evolution of aggressiveness in a local plant pathogen population. This time-scale effect is consistent with theoretical results suggesting that counter-selection may occur during interepidemic periods (van den Berg *et al*., 2011) and with experimental results obtained with other plant pathogens suggesting a functional, adaptive compromise between the parasitic and saprophytic survival stages (Abang *et al.*, 2006; Sommerhalder *et al*., 2011; Laine & Barrès, 2013; Susi & Laine, 2013; Pasco *et al*., 2015). Caffier *et al*. (2016) showed a slow erosion of quantitative resistance to *Venturia inaequalis,* conferred by a single isolate-specific QTL, in apple trees. Aggressive isolates were selected, but no change in aggressiveness or in the frequency of the most aggressive isolates was detected over an eight-year period. A trade-off between capacity to overcome the QTL and capacity to keep in *V. inaequalis* populations over years was suggested; it may be due to counter-selection during the interepidemic period (fewer opportunities to reproduce sexually due to the lower density of lesions on the leaves). Similar experiments assessing the erosion of moderate and high levels of resistance to *Z. tritici,* taking into account the deployment of the cultivars at the landscape scale, would be relevant, as already done for *P. triticina* (Papaïx *et al*., 2011; 2014). It should be noted that assessments of the aggressiveness of the isolates on the allopatric host (cv. Apache) did not highlight this trade-off. Similar apparently inconsistent results obtained for cultivar mixtures were interpreted as an expression of disruptive selection affecting the evolution of *Z. tritici* populations (Mundt *et al.*, 1999). The aggressiveness of the pathogen population *per se* should therefore not be considered independently of the nature and level of host resistance.

The strength of the difference in the evolution of aggressiveness at the intra- and interepidemic periods is probably determined by the balance between processes leading to selection over the course of a single annual epidemic and processes hampering interepidemic transmission. It is clearly challenging to assess this balance in field conditions. In our study area, during a growing season conducive to disease (2009-2010), the pathogen probably completed six asexual infection cycles. We therefore compared the effects of these six asexual reproduction cycles with those of six sexual reproduction cycles over the six-year duration of the whole experiment.

As no functional trade-off between asexual and sexual reproduction was found at the plant scale in previous experiments (Suffert *et al*., 2015; 2016), we suggest that the counter-selection observed during the interepidemic period results principally from an Allee effect. Indeed, more aggressive *Z. tritici* isolates (Cf-2010) were collected from the upper leaves, on which disease severity was generally lower than on the lower parts of the plants (wheat basal leaves and stems) then left in the field as crop debris after harvest. A low pathogen density on the upper part of the plants decreases the likelihood of isolates of compatible mating types meeting. The likelihood of the isolates present on upper leaves reproducing sexually is further reduced by the frequent removal of the upper parts of the plants from the field during harvesting, whereas the lower parts of the plants tend to be left behind. Sexual reproduction in *Z. tritici* populations is even more intense on the basal parts of wheat plants when the plants considered are old and disease severity is high (Eriksen & Munk, 2003), as indicated by the positive correlation between winter disease severity (reflecting pathogen density on lower leaf layers) and the subsequent amount of primary inoculum (reflecting the intensity of sexual reproduction). These processes probably provide the best explanation for the low interannual transmissibility of the most aggressive pathogen isolates selected during the annual epidemic for their ability to propagate clonally, and for the difference in the evolution of aggressiveness in the pathogen population over the intra- and interannual scales.

The results of our study call for more thorough investigations of the quantitative balance between epidemiological processes leading to a trade-off relationship in the evolution of aggressiveness during the intra- and interepidemic periods, for improving the deployment of host resistance in a landscape over several years. This issue is particularly important for pathogens of annual crops that, like *Z. tritici,* have a dual reproduction cycle, including a saprophytic survival stage on crop debris (e.g. *Rhynchosporium secalis*, Abang *et al.*, 2006; *Phaeosphaeria nodorum*, Sommerhalder *et al*., 2011). This issue is also important for strict biotrophic pathogens of annual or perennial crops, such as rusts (e.g. *Melampsora larici-populaina*, Pernaci, 2015; *P. triticina*, Soubeyrand *et al*., 2017), for which alternative hosts or volunteers act as a green bridge during the interepidemic period. The epidemiological processes involved depend on the biology of the pathogen, climatic conditions and agronomic context. Changes in the management of crop debris and volunteers in crop systems, such as the development of simplified cultivation practices, may account for the past or future evolution of aggressiveness in pathogens.

## ACKNOWLEDGMENTS

We thank Nathalie Galet, Christian Lepoulennec (INRA BIOGER) and Christophe Montagnier (INRA Experimental Unit, Thiverval-Grignon) for technical assistance during this long-term experiment and Thierry Marcel (INRA BIOGER) for helpful discussions on the determinism of wheat resistance to Septoria tritici blotch. We thank FranceAgriMer and ARVALIS-Institut du Végétal for providing data concerning the proportion of wheat cultivars in France and in the Yvelines in particular. We thank Benoit Moury and the second anonymous reviewer, both members of PCI Evol Biol (Peer Community in Evolutionary Biology; https://evolbiol.peercommunityin.org/), for their insightful comments on the manuscript. We thank J. Sappa for her help correcting our English. This study was supported by a grant running from 2007 to 2013 under the European Union Seventh Framework Program (Grant Agreement no. 261752, PLANTFOODSEC project), a 2015-2019 grant from the European Union Horizon 2020 program (Grant Agreement no. 634179, EMPHASIS project), and a 2014-2016 grant from LabEx BASC (SEPTOVAR project) as part of the “Investissements d’Avenir” program (ANR-11-LABX-0034).

## Supplementary Material

**Table S1.** *Zymoseptoria tritici* subpopulations (2 × 2 × 15 isolates) sampled in a wheat monoculture plot (Grignon, France).

| Ci-2009 |  |  | Cf-2010 |  |  |
| --- | --- | --- | --- | --- | --- |
| Name | Code | Date of collection | Name | Code | Date of collection |
| INRA09-FS0729 | I01 | 24 November 2009 | INRA09-FS01000 | I16 | 12 July 2010 |
|  |  |  | INRA09-FS01002 | I22 |  |
| INRA09-FS0731 | I06 | 30 November 2009 | INRA09-FS01003 | I23 |  |
| INRA09-FS0732 | I07 |  | INRA09-FS01006 | I24 |  |
|  |  |  | INRA09-FS01008 | I25 |  |
| INRA09-FS0798 | I02 | 8 December 2009 | INRA09-FS01013 | I17 |  |
| INRA09-FS0799 | I08 |  | INRA09-FS01015 | I26 |  |
| INRA09-FS0800 | I09 |  | INRA09-FS01018 | I18 |  |
| INRA09-FS0802 | I03 |  | INRA09-FS01019 | I27 |  |
| INRA09-FS0803 | I10 |  | INRA09-FS01021 | I19 |  |
| INRA09-FS0805 | I11 |  | INRA09-FS01022 | I28 |  |
| INRA09-FS0806 | I12 |  | INRA09-FS01023 | I29 |  |
| INRA09-FS0808 | I04 |  | INRA09-FS01024 | I20 |  |
| INRA09-FS0809 | I13 |  | INRA09-FS01025 | I21 |  |
| INRA09-FS0811 | I14 |  | INRA09-FS01026 | I30 |  |
| INRA09-FS0813 | I05 |  |  |  |  |
| INRA09-FS0814 | I15 |  |  |  |  |
| Ai-2009 |  |  | Af-2015 |  |  |
| Name | Code | Date of collection | Name | Code | Date of collection |
| INRA09-FS0402 | I31 | 9 October 2009 | INRA09-FS0265 | I46 | 7 September 2015 |
| INRA09-FS0406 | I32 |  | INRA09-FS0266 | I47 |  |
| INRA09-FS0410 | I33 |  | INRA09-FS0267 | I48 |  |
| INRA09-FS0411 | I34 |  | INRA09-FS0268 | I49 |  |
| INRA09-FS0414 | I35 |  | INRA09-FS0269 | I50 |  |
| INRA09-FS0417 | I36 |  | INRA09-FS0270 | I51 |  |
| INRA09-FS0420 | I37 |  | INRA09-FS0271 | I52 |  |
| INRA09-FS0421 | I38 |  | INRA09-FS0278 | I53 |  |
| INRA09-FS0423 | I39 |  |  |  |  |
| INRA09-FS0425 | I40 |  | INRA09-FS0272 | I54 | 13 October 2015 |
| INRA09-FS0439 | I41 |  | INRA09-FS0273 | I55 |  |
| INRA09-FS0434 | I42 |  | INRA09-FS0274 | I56 |  |
| INRA09-FS0438 | I43 |  | INRA09-FS0275 | I57 |  |
| INRA09-FS0444 | I44 |  | INRA09-FS0276 | I58 |  |
| INRA09-FS0452 | I45 |  | INRA09-FS0277 | I59 |  |
|  |  |  | INRA09-FS0280 | I60 |  |

